# Case-control vs. case-only estimates of gene-environment interactions with common and misclassified clinical diagosis

**DOI:** 10.1101/760322

**Authors:** Iryna Lobach, Ying Sheng, Siarhei Lobach, Lydia Zablotska, Chiung-Yu Huang

## Abstract

Genetic studies provide valuable information to assess if the effect of genetic variants varies by the non-genetic (“environmental”) variables, what is traditionally defined to be gene-environment interaction. A common complication is that multiple disease states present with the same set of symptoms, and hence share the clinical diagnosis. Because 1) disease states might have distinct genetic bases; and 2) frequencies of the disease states within the clinical diagnosis vary by the environmental variables, analyses of association with the clinical diagnosis as an outcome variable might result in false positive or false negative findings. We develop estimates for assessment of GxE in a case-only study and compare the case-control and case-only estimates. We report extensive simulation studies that evaluate empirical properties of the estimates and show the application to a study of Alzheimer’s disease.

## INTRODUCTION

Numerous genome-wide association studies (GWAS) have been conducted to estimate how the effect of genetic variants varies by non-genetic (environmental) variables, what is traditionally referred to as gene-environment interactions (GxE). A major and commonly overseen complication is that multiple distinct pathophysiologic mechanisms might present with the same set of symptoms and hence the same clinical diagnosis. Frequencies of the disease states within the clinically diagnosed set often vary by the environmental variables, such as age, race/ethnicity. The pathophysiologic mechanisms underlying the disease states might have distinct genetic bases. Hence the analyses with the clinical diagnosis as an outcome variable might miss important associations or result in spurious findings (Carroll et al, 2016).

Our study is motivated by the setting of Alzheimer’s disease (AD) where approximately 30% of patients clinically diagnosed with AD do not have evidence of amyloid deposition as measured by positron emission tomography (PET) (Ossenkoppele et al, 2015). Hence two disease states – symptoms of AD with amyloid evidence and symptoms of AD with no amyloid evidence – are within the clinical diagnosis of AD. We define the disease state of AD with amyloid evidence to be the disease state of interest (Potter and Wisniewski, 2012). Frequencies of the disease states within the clinical diagnosis are estimated to vary by age and Apolipoprotein (ApoE) *ε*4 status (Ossenkoppele et al, 2015). Interestingly, ApoE *ε*4 status is the most potent genetic factor found thus far. Genetic studies that define the clinical diagnosis as an outcome variable found that risk of AD increases and the age at onset decreases with the number of ApoE *ε*4 alleles (Corder et al, 1993; Farrer et al, 1997), thus suggesting a GxE. Because the disease states of AD with and without the amyloid evidence might have distinct genetic bases and the mechanism of ApoE *ε*4 action might be relevant to the amyloid deposition, we are interested to assess the GxE in the relationship to the disease state of AD underlined by the amyloid evidence.

Our previous studies showed both empirically and theoretically that ignoring heterogeneity of AD diagnosis can lead to severely biased estimates of GxE (Lobach et et al, 2018; Lobach et al, 2019).

We are interested to estimate the GxE from the set of clinically diagnosed cases only, assuming that G and E are distributed independently in the population. This interest is supported by the prior statistical literature showing that when the genotype and environment are distributed independently in the population and when the disease is rare, the GxE can be estimated from a case-only study more efficiently than from a case-control study (Piegorsch et al, 1994). This result, however, is not applicable to our setting both because the disease state and the clinical diagnoses are common, and because the clinical diagnosis is not a *surrogate* of the disease state. We, therefore, are interested to derive what types of GxE can be estimated from the set of clinically diagnosed cases and compare variability of case-control vs. case-only estimates.

Our paper proceeds as follows. We first describe the setting and derivations in the Case-control vs. case-only Estimates section. Next, we evaluate the estimates in empirical studies and describe the setting and results in the Simulation Studies section. The application of the methods is then shown on a large-scale study of Alzheimer’s disease. We conclude the paper with brief discussion.

## Materials and Methods

### Case-control vs. case-only estimates

We consider a study consisting of *n*_1_ cases with a clinical diagnosis and *n*_0_ controls. The data are collected using retrospective sampling scheme, i.e. cases are collected from the population of clinically diagnosed cases and controls are collected from the population of the clinically diagnosed controls. Suppose that what measured is a set of genotypes *G* and environmental variables *E* that are distributed independently in the population. We define the observed clinical diagnosis be *D*^*CL*^ = {0,1} and the true disease state to be *D* = {0,1}. We let 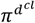 be frequency of the clinical diagnosis in the population, and *π*^*d*^ -the frequency of the disease state in the population. We define frequencies of the disease state of interest within the clinical diagnosis as *pr*(*D* = 1|*D*^*CL*^ = 1, *G* = *g, E* = *e*) = *τ*(*g, e*). For clarity of the presentation we will assume that the set of controls is homogeneous, i.e. *pr*(*D* = 0|*D*^*CL*^ = 0, *E* = *e, G* = *g*) = 1. For clarity of the presentation we suppose that all variables are binary.

We define *Q*(g|θ) be the distribution of genotype in the population according to Hardy-Weinberg Equilibrium.

We define frequencies of the genotype and environment within the clinical diagnosis and the disease states to be 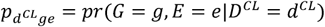 and *π*_*dg*e_ = *pr*(*G* = *g, E* = *e*|*D* = *d*).

We are interested to assess GxE. The traditional analyses are based on the logistic regression model, where GxE is a multiplicative interaction capturing the deviation from the sum of main effects of G and E. We hence start by considering a logistic regression models with the disease state of interest as an outcome variable, where the interaction term is of the primary interest. In the context of this study we are not interested in estimating the main effects and hence the risk model itself, we are just aiming to assess if the data provides sufficient evidence for an interactive effect. Hence consider a model

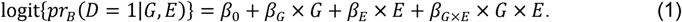

The disease states, however, are not observed directly, instead what is measured is a clinical diagnosis *D*^*CL*^ defined based on the set of observed symptoms. Hence the observed data allows us to estimate an interaction term from the following model

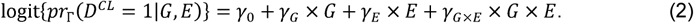

If the clinical diagnosis is a *surrogate* of the disease state of interest, i.e. pr(*D*^*CL*^ = *d*^*cl*^|*D* = *d, G* = *g, E* = *e*) = pr(*D*^*CL*^ = *d*^*cl*^|*D* = *d*), then 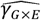 is a consistent estimate of *β*_*G*×*E*_. In this setting 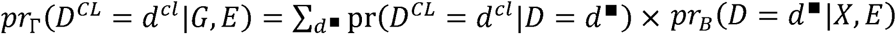, hence if there is no relationship between (*G, E*) and *D*, neither there is one between (*G, E*) and *D*^*C*L^. Otherwise, the probabilities of the clinical diagnosis are weighted sums of frequencies of the true diagnosis, 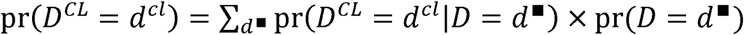, and 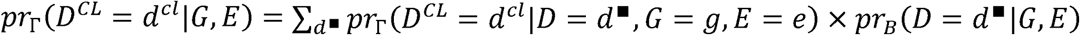. Then if there is no relationship between (*G, E*) and *D*, there might be the relationship between (*G, E*) and *D*^*CL*^.

The seminal work by Piegorsch er al (1994) developed a multiplicative interaction for a rare disease assuming independence between *G* and *E*, i.e. for *β*_*G*×*E*_, as follows. An odds ratio (OR) for *E* in *G* = 0 is then 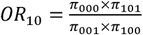, for *G* in *E* = 0 is 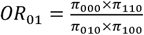 and for *G* = 1, *E* = 1 vs. *G* = 0, *E* = 0 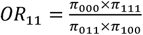. A multiplicative interaction is then 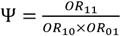 and can be estimated in a case-control study as

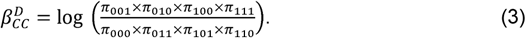

The case-only estimate is

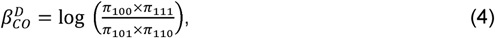

which is not applicable to our study for two reasons. First, because the disease states and the clinical diagnosis are not rare. Second, because some of the clinically diagnosed cases are misdiagnosed controls.

Because 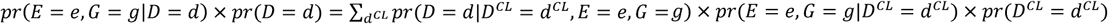, the GxE interaction in (1), i.e. see (3), becomes

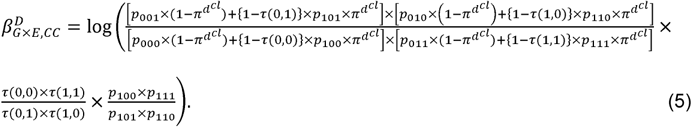

The case-control GxE estimate (5) cannot be seamlessly reduced to a case-only estimate following the arguments of Piegorsh et al (1994) mainly because the disease state and the clinical diagnosis are not rare.

Hence, we are interested to derive other estimates that characterize how the effect of genotype varies by the environment and that can be estimated in a set of clinically diagnosed cases. We aim to derive estimates of GxE that are necessary and might not be sufficient for evaluating whether or not GxE is present.

It can be easily seen that the environment-specific odds of genotype among cases with the disease state of interest is

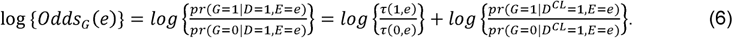

Recall that *θ*= *pr*(*G*= 1) and then the environment-specific risk ratio attributable to genotype is

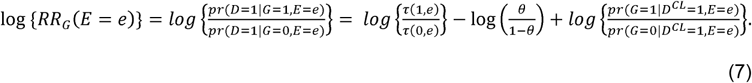

From the statistical literature, including the study by Piegorsh et al (1994), we know that a case-only estimate for GxE can be obtained from regressing the environment on the genotype within the set of cases. i.e. a case-only estimate of GxE is the coefficient *α*_*E*_ from the logistic model

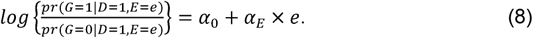

The coefficient *α*_*E*_ defines GxE on the relative risk scale. Then this model in combination with (6) arrives at

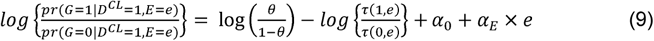

The analog of model (8) with the clinical diagnosis as an outcome variable is

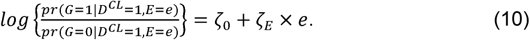

Hence the case-only GxE coefficient *α*_*E*_ can be estimated from the set of clinically diagnosed cases as

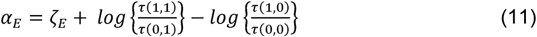

We assume that frequency of genotype, *θ*, and frequencies of the disease states within the clinical diagnosis, *τ*(*g, e*), are known. We define *c*_*e*_(*g*) = #(*G* = *g*|*D*^*CL*^ = 1, *E* = *e*). Then variance of the risk ratios (7) and (11) is

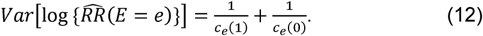

Similarly, variance of the odds (6) is

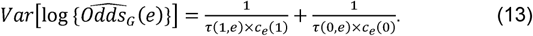

### Remarks

1. If the clinical diagnosis is rare in the population, i.e.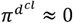, then we can see from (5) that the estimate with the clinical diagnosis as an outcome variable is not an unbiased estimate of the GxE for the disease state of interest. That is for rare diseases ignoring misclassification of the diagnosis does not lead to unbiased estimates of GxE.
2. In Alzheimer’s disease study that motivated this work, frequencies of the disease states within the clinical diagnosis are a function of both the genotype and the environment, i.e. *pr*(*D*= 1|*D*^*CL*^ = 1, *G* = *g, E* = *e*) = *τ*(*g, e*).
3. If the frequency is only a function of the genotype, i.e. *τ*(*g, e*) = *τ*(*g*), or the environment, i.e. *τ*(*g, e*) = *τ*(*e*); then *α*_*E*_ = *ζ*_*E*_ but the estimate (5) does not reduce to the estimate (3). This setting occurs when the main effect of the environment or genotype is null.
4. We’ve derived the case-control GxE estimate (5) and proposed three measures to assess how the effect of genotype varies by the environment from a set of clinically diagnosed cases: environment-specific odds of genotype within the subset with the disease state of interest 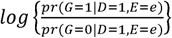 (6), environment-specific risk ratio attributable to the genotype 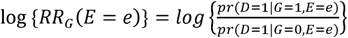(7), and the coefficient *α*_*E*_ (9).
5. All the risk ratios and odds ratios that we discussed can be easily redefined to be genotype-specific, for example the genotype-specific odds of the environment

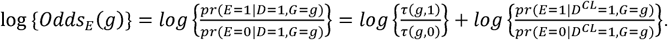

## Simulation Studies

We conduct a series of simulation studies to asses bias and false discovery rates of the estimates that ignore presence of the nuisance disease state within the clinical diagnosis, i.e. estimates (3) and (4) with the clinical diagnosis in place of the disease state. We also assess performance of the odds (6), the risk ratio (7), and the estimate (11); and their variances (12), (13).

In all settings we simulate 500 samples of with *n*_0_ clinically diagnosed controls and *n*_1_ clinically diagnosed cases. We let the genetic variant be Bernoulli(0.10) and the environmental variable be Bernoulli(0.14). We then simulate the disease state *D* according to the risk model (1) with values of the coefficients *β*_0_, *β*_*G*_, *β*_*E*_, *β*_*G*×*E*_ that vary. We next simulate the clinical diagnosis according to frequencies of the nuisance disease state within the clinical diagnosis with *pr*(*D* = 1|*D*^*CL*^ = 1, *E* = *e, G* = *g*) = *τ*(*e, g*).

We let *β*_0_ = −0.5, *β*_*G*_ = log(1), log(2), log(5), *β*_*E*_ = log(2). We let {*pr*(*D* = 1|*D*^*CL*^ = 1, *E* = 0), *pr* (*D* = 1|*D*^*CL*^ = 1, *E* = 1)} be {0.64, 0.94}; {0.64, 0.84}; {0.84, 0.94}, i.e. we vary both the proportion of the nuisance disease states within the clinically diagnosed set of cases and the difference in the proportion by *E*.

### Setting 1: Null effect, i.e. *β*_*G*×*E*_ = 0

We first consider a setting when there is no GxE, i.e. *β*_*G*×*E*_ = 0. Shown in **Table 1A** are biases, SDs in the estimates, as well as false discovery rates (FDR). We note that the FDR in the estimates that use the clinical diagnosis as an outcome variable ranges between 0.06 and 0.40 across the settings we considered. Hence these inferences can be substantially inflated. The estimates with the disease state as an outcome variable, i.e. (5) and (11), are nearly unbiased. The case-only estimates tend to have variability that is less or equal to the variability of the case-control estimates, with relative efficiency varying from 1 to 2.4. Shown in **Table 1B** are the odds (6) and risk ratio (7). The estimates are nearly unbiased with empirical variability that is approximately the same as the theoretical variance (12), as shown in **Table 1C**.

**Table 1A:** Biases and standard deviations (SD) of case-control and case-only estimates when the clinical diagnosis is used in place of the disease state ((3) and (4)) and when the outcome variable is the disease state ((5) and (11)). We simulated 500 datasets with 3,000 clinically diagnosed cases and 3,000 clinically diagnosed controls where the disease state is simulated according to model (1) with coefficients *β*_*G*×*E*_ = log (0), *β*_0_ = −0.5, *β*_*G*_ = log(1), log(2), log(5), *β*_*E*_ = log(3) and the clinical diagnosis is then simulated according to frequencies *pr*(*D* = 1|*D*^*CL*^ = 1, *G* = *g, E* = *e*).

| $pr(D = 1\vert D^{CL} = 1, G = 0)$ | $pr(D = 1\vert D^{CL} = 1, G = 1)$ | $\beta_G$ | Clinical diagnosis as an outcome variable | | | | | | Disease state is the outcome variable | | | | |
| --- | --- | --- | --- | --- | --- | --- | --- | --- | --- | --- | --- | --- | --- |
|  |  |  | Case-control estimate |  |  | Case-only estimate |  |  | Case-control estimate |  | Case-only estimate |  |  |
|  |  |  | Bias | SD | False positive rate | Bias | SD | False positive rate | Bias | SD | True value | Bias | SD |
| 0.64 | 0.94 | Log(1)=0 | 0.21 | 0.15 | 0.29 | 0.20 | 0.12 | 0.40 | -0.12 | 0.13 | 0.00 | -0.0002 | 0.12 |
|  |  | Log(2)=0.69 | 0.21 | 0.16 | 0.30 | -0.21 | 0.12 | 0.14 | -0.11 | 0.12 | -0.12 | 0.01 | 0.12 |
|  |  | Log(5)=1.61 | 0.21 | 0.18 | 0.22 | -0.03 | 0.10 | 0.07 | 0.09 | 0.11 | -0.25 | 0.05 | 0.10 |
| 0.64 | 0.84 | Log(1)=0 | 0.14 | 0.14 | 0.15 | 0.14 | 0.12 | 0.22 | -0.08 | 0.12 | 0.00 | -0.0007 | 0.12 |
|  |  | Log(2)=0.69 | 0.15 | 0.16 | 0.16 | 0.03 | 0.11 | 0.09 | -0.02 | 0.10 | -0.13 | 0.02 | 0.11 |
|  |  | Log(5)=1.61 | 0.17 | 0.27 | 0.08 | -0.10 | 0.10 | 0.20 | 0.12 | 0.16 | -0.24 | -0.004 | 0.10 |
| 0.84 | 0.94 | Log(1)=0 | 0.07 | 0.15 | 0.08 | 0.07 | 0.11 | 0.09 | -0.03 | 0.14 | 0.008 | 0.003 | 0.11 |
|  |  | Log(2)=0.69 | 0.07 | 0.16 | 0.09 | -0.04 | 0.11 | 0.09 | -0.002 | 0.14 | 0.02 | -0.11 | 0.11 |
|  |  | Log(5)=1.61 | 0.08 | 0.20 | 0.06 | -0.17 | 0.09 | 0.47 | 0.07 | 0.14 | -0.24 | 0.001 | 0.09 |

**Table 1B:**
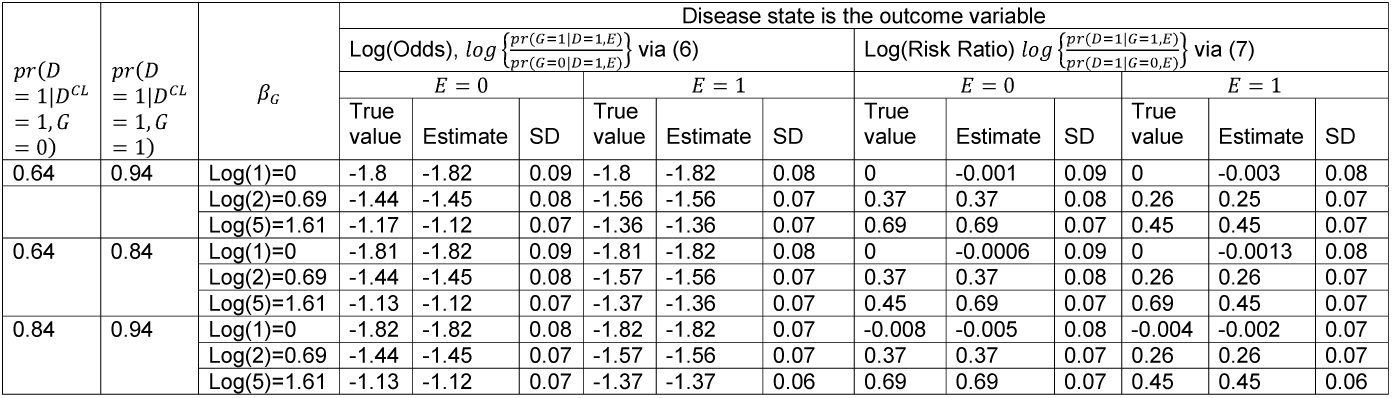
True values, empirical averages and standard deviations (SD) of case-only odds (6) and risk ratio (7) when the outcome variable is the disease state. We simulated 500 datasets with 3,000 clinically diagnosed cases and 3,000 clinically diagnosed controls where the disease state is simulated according to model (1) with coefficients *β*_*G*×*E*_ = log (0), *β*_0_ = −0.5, *β*_*G*_ = log(1), log(2), log(5), *β*_*E*_ = log(3) and the clinical diagnosis is then simulated according to frequencies *pr*(*D* = 1|*D*^*CL*^ = 1, *G* = *g, E* = *e*).

**Table 1C:** Empirical averages of the theoretical standard deviations (SD) as described in (12). We simulated 500 datasets with 3,000 clinically diagnosed cases and 3,000 clinically diagnosed controls where the disease state is simulated according to model (1) with coefficients *β*_*G*×*E*_ = log (0), *β*_0_, =−0.5, *β*_*G*_ = log (1) log (2), log (5), *β*_*E*_ = log (3) and the clinical diagnosis is then simulated according to frequencies *pr(D* = 1|*D*^*CL*^ = 1, *G* = *g, E* = *e*).

| $pr(D = 1\vert D^{CL} = 1, G = 0)$ | $pr(D = 1\vert D^{CL} = 1, G = 1)$ | $\beta_G$ | Disease state is the outcome variable | |
| --- | --- | --- | --- | --- |
| | | | $E = 0$ | $E = 1$ |
|  |  |  | Theoretical SD |  |
| 0.64 | 0.94 | Log(1)=0 | 0.09 | 0.08 |
|  |  | Log(2)=0.69 | 0.08 | 0.07 |
|  |  | Log(5)=1.61 | 0.07 | 0.07 |
| 0.64 | 0.84 | Log(1)=0 | 0.09 | 0.08 |
|  |  | Log(2)=0.69 | 0.08 | 0.07 |
|  |  | Log(5)=1.61 | 0.07 | 0.07 |
| 0.84 | 0.94 | Log(1)=0 | 0.08 | 0.07 |
|  |  | Log(2)=0.69 | 0.07 | 0.07 |
|  |  | Log(5)=1.61 | 0.07 | 0.06 |

### Setting 2: *β*_*G*×*E*_ = log(3)

We then consider a setting where *β*_*G*×*E*_ = log (3). Estimates shown **Table 2A** can be substantially biased when the clinical diagnosis is used in place of the outcome variable, while the bias in nearly removed when the disease state is the outcome. The case-only estimates tend to have variability that is smaller than the variability of the case-control estimates with relative efficiencies varying between 1 and 1.7. Risk ratio estimates in **Table 2B** are nearly unbiased with empirical variability that is close to the theoretical as shown in **Table 2C**.

**Table 2A:**
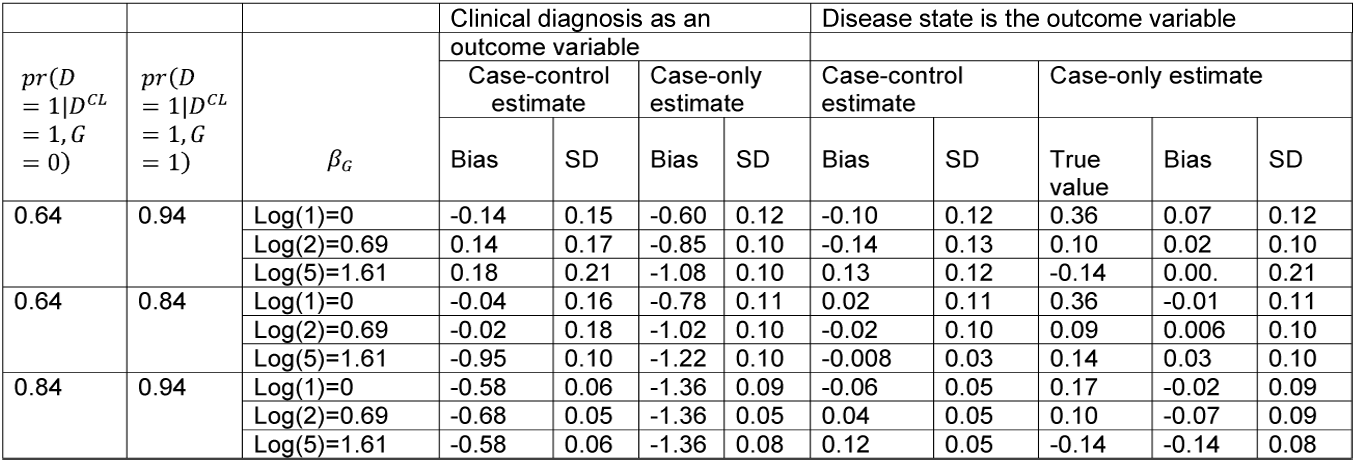
Biases and standard deviations (SD) of case-control and case-only estimates when the clinical diagnosis is used in place of the disease state ((3) and (4)) and when the outcome variable is the disease state ((5) and (11)). We simulated 500 datasets with 3,000 clinically diagnosed cases and 3,000 clinically diagnosed controls where the disease state is simulated according to model (1) with coefficients *β*_*G*×*E*_ = log (3), *β*_0_ = −0.5, *β*_*G*_ = log(1), log(2), log(5), *β*_*E*_ = log(3) and the clinical diagnosis is then simulated according to frequencies *pr*(*D* = 1|*D*^*CL*^ = 1, *G* = *g, E* = *e*).

**Table 2B:**
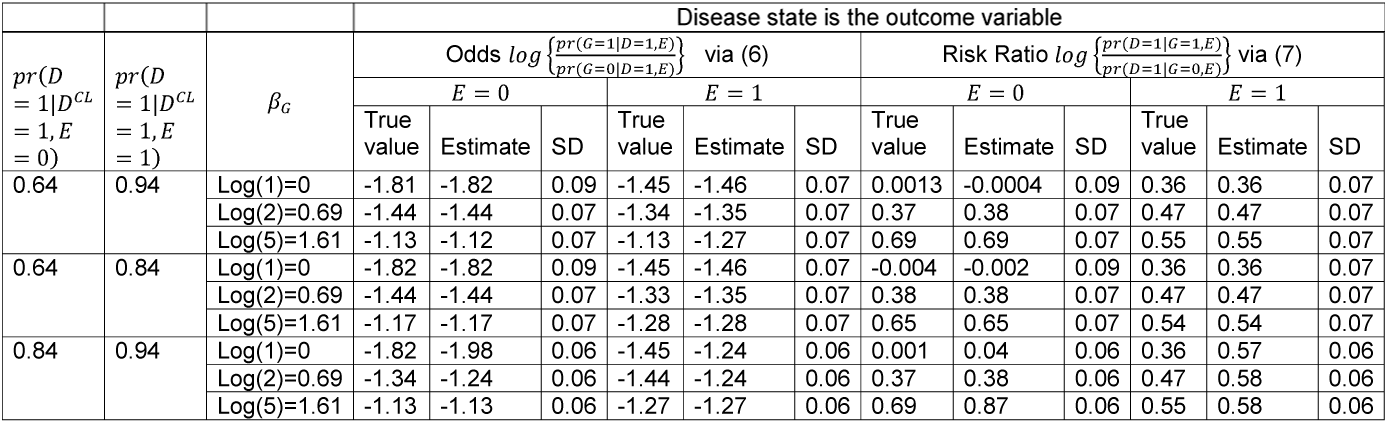
True values, empirical averages and standard deviations (SD) of case-only odds (6) and risk ratio (7) when the outcome variable is the disease state. We simulated 500 datasets with 3,000 clinically diagnosed cases and 3,000 clinically diagnosed controls where the disease state is simulated according to model (1) with coefficients *β*_*G*×*E*_ = log (3), *β*_0_ = ×0.5, *β*_*G*_ = log(1), log(2), log(5), *β*_*E*_ = log(3) and the clinical diagnosis is then simulated according to frequencies *pr*(*D* = 1|*D*^*CL*^ = 1, *G* = *g, E* = *e*).

**Table 2C:** Empirical averages of the theoretical standard deviations (SD) as described in (12). We simulated 500 datasets with 3,000 clinically diagnosed cases and 3,000 clinically diagnosed controls where the disease state is simulated according to model (1) with coefficients *β*_*G*×*E*_ = log (3), *β*_0_ = −0.5, *β*_*G*_ = log (1), log (2), log (5), *β*_*E*_ = log (3) and the clinical diagnosis is then simulated according to frequencies *pr*(*D* = 1|*D*^*CL*^ = 1, *G* = *g, E* = *e*).

| $pr(D = 1\vert D^{CL} = 1, G = 0)$ | $pr(D = 1\vert D^{CL} = 1, G = 1)$ | $\beta_G$ | Disease state is the outcome variable | |
| --- | --- | --- | --- | --- |
| | | | $E = 0$ | $E = 1$ |
|  |  |  | Theoretical SD |  |
| 0.64 | 0.94 | Log(1)=0 | 0.09 | 0.09 |
|  |  | Log(2)=0.69 | 0.07 | 0.07 |
|  |  | Log(5)=1.61 | 0.07 | 0.07 |
| 0.64 | 0.84 | Log(1)=0 | 0.09 | 0.07 |
|  |  | Log(2)=0.69 | 0.07 | 0.07 |
|  |  | Log(5)=1.61 | 0.07 | 0.06 |
| 0.84 | 0.94 | Log(1)=0 | 0.06 | 0.06 |
|  |  | Log(2)=0.69 | 0.06 | 0.06 |
|  |  | Log(5)=1.61 | 0.07 | 0.06 |

## Alzheimer’s disease study

We applied the proposed analyses to a dataset collected as part of the Alzheimer’s Disease Genetics Consortium (Naj et al, 2011). The data consists of 1,245 controls and 2,785 cases. The average age (SD) of cases and controls are 72.1 (9.1) and 70.9 (8.8) years, respectively. Among cases, 1,458 (52.4%) are men; among controls, 678 (63.9%) are men. At least one ApoE *ε*4 allele is present in (64.5%) of cases and 365 (29.1%) of controls.

We are considering late-onset AD, hence focus on the subpopulation aged 65 years or older.

Recent biomarker studies (Ossenkoppele et al, 2015) estimate that 95% of ApoE *ε*4 carriers and 78% of ApoE *ε*4 non-carriers aged 65-75 diagnosed with AD have evidence of amyloid deposition. Similarly, 90% of ApoE *ε*4 carriers and 63% of ApoE *ε*4 non-carriers aged 75+ clinically diagnosed with AD have evidence of amyloid deposition. We assume that 1 in 10 is diagnosed with AD (https://www.alzheimers.net/resources/alzheimers-statistics/). We will perform sensitivity analyses by assuming these rates and also varying the rates to see how the estimates of ApoE *ε*4-by-age interaction might change.

In **Table 3** we present various estimates of how the effect of ApoE *ε*4 varies by age with 95% Confidence Intervals (CI) obtained based on 1,000 permutations. We first note that the case-control estimate with the clinical diagnosis as an outcome variable is not statistically significant (−0.08; 95% CI (−0.30, 0.29)), while the case-control estimate with the disease state as an outcome variable is statistically significant (−0.69; 95% CI (−0.76, -0.58)). Across all the settings we considered, length of the 95% CI for the case-only estimates tend to be shorter than the length for the case-control estimate. Setting 1 presented in **Table 3** corresponds to the estimates of the disease states obtained in Ossenkoppele et al, 2015; while Settings 2-4 are slight deviations. We note that the estimates and their 95% CIs are similar across all the settings showing robustness of the conclusions to the slight derivations in the estimates of the disease states within the clinical diagnosis.

**Table 3:** Estimates (95% Confidence Intervals) of how the effect of ApoE *ε*4 varies by age in the Alzheimer’s disease study. Setting 1 is as estimated in the literature. The other Settings are slight deviations from the Setting 1 for analyses of sensitivity.

| Frequencies of the AD disease state with amyloid evidence within the clinical | Setting 1 | Setting 2 | Setting 3 | Setting 4 |
| --- | --- | --- | --- | --- |
| diagnosis by age |  |  |  |  |
| $pr(D = 1\vert D^{CL} = 1, Age = 65 - 75, \varepsilon 4+) )$ | 95% | 95% | 90% | 100% |
| $pr(D = 1\vert D^{CL} = 1, Age = 75+, \varepsilon 4+) )$ | 90% | 90% | 85% | 95% |
| $pr(D = 1\vert D^{CL} = 1, Age = 65 - 75, \varepsilon 4- )$ | 78% | 78% | 73% | 83% |
| $pr(D = 1\vert D^{CL} = 1, Age = 75+, \varepsilon 4- )$ | 63% | 63% | 58% | 68% |
| $pr(D^{CL} = 1)$ | 10% | 7% | 10% | 10% |
| Clinical diagnosis is the outcome variable |  |  |  |  |
| Case-control estimate | -0.08 (-0.30, 0.29) |  |  |  |
| Case-only estimate | **-0.69** (-0.76, -0.58) |  |  |  |
| Disease state is the outcome variable |  |  |  |  |
| Case-control estimate, (5) | **0.89** (0.83, 0.95) | **0.88** (0.82, 0.94) | **0.91** (0.84, 0.96) | **0.87** (0.81, 0.93) |
| Case-only estimate, (11) | **0.83** (0.73, 0.93) | **0.82** (0.74, 0.92) | **0.84** (0.75, 0.93) | **0.82** (0.72, 0.90) |
| Odds $\log \left\{ \frac{pr(\varepsilon 4+\vert D=1, Age=65-75)}{pr(\varepsilon 4-\vert D=1, Age=65-75)} \right\}$, (6) | **0.63** (0.58, 0.69) | **0.63** (0.58, 0.70) | **0.65** (0.59, 0.70) | **0.62** (0.57, 0.68) |
| Odds $\log \left\{ \frac{pr(\varepsilon 4+\vert D=1, Age=75+)}{pr(\varepsilon 4-\vert D=1, Age=75+)} \right\}$, (6) | **0.12** (0.04, 0.20) | **0.12** (0.04, 0.20) | **0.15** (0.08, 0.23) | **0.10** (0.03, 0.18) |
| Risk Ratio $\log \left\{ \frac{pr(D=1\vert \varepsilon 4+, Age=65-75)}{pr(D=1\vert \varepsilon 4-, Age=65-75)} \right\}$, (7) | **2.4** (2.4, 2.5) | **2.4** (2.4, 2.5) | **2.5** (2.4, 2.5) | **2.4** (2.3, 2.5) |
| Risk Ratio $\log \left\{ \frac{pr(D=1\vert \varepsilon 4+, Age=75+)}{pr(D=1\vert \varepsilon 4-, Age=75+)} \right\}$, (7) | **2.3** (1.9, 2.0) | **1.9** (1.9, 2.0) | **2** (1.9, 2.0) | **1.9** (1.8, 2.0) |

## Discussion

We’ve derived case-control and case-only estimates of GxE with respect to the AD disease state underlined by amyloid deposition in the setting when the disease state is not measured, instead frequencies of the disease state within the clinical diagnosis are estimated in a reliability study. We also evaluated bias and false discovery rates for when the misclassification of the clinical diagnosis is ignored. The setting we consider is unique in that the disease states and the clinical diagnosis are common and that rates of the disease state of interest within the clinical diagnosis vary by G and by E.

The development of our study is motivated by the need to assess presence of GxE, as opposed to estimating all parameters in a risk model, e.g. (1). We are interested to estimate the degree to which the effect of genotype varies by the “environment”, such as age, sex, education, race/ethnicity, etc. The setting that we’ve developed offers an advantage of not having to rely on the estimates of main effects.

In simulation experiments we showed that ignoring misclassification of the clinical diagnosis can result in substantial inflation of false positive rates in GxE. Similarly to the original study by Piegorsch et al (1994), we note that the variability of case-only estimates is generally smaller than the variability of the case-control estimates.

The derivations that we’ve developed rely on estimates of the population frequencies of the disease states of interest within the clinical diagnosis that vary by G and E. These estimates are often available. We advocate for sensitivity analyses by assuming the estimates obtained in reliability studies and varying the values slightly to see if the GxE estimates change.

While our study is motivated by the setting of Alzheimer’s disease, the general development is applicable to other complex diseases and other settings, e.g. analyses of association in the context of the electronic health records, or other types of genetic data, e.g. gene expression. For example, studies of diabetes (Manchia et al, 2013) and analyses of electronic health records (Hubbard et al, 2017).

## ACKNOLEDEGMENTS

Dr. Lobach is supported by 5R21AG043710-02.

Genotyping is performed by Alzheimer’s Disease Genetics Consortium (ADGC), U01 AG032984, RC2 AG036528. Phenotypic collection is coordinated by the National Alzheimer’s Coordinating Center (NACC), U01 AG016976

Samples from the National Cell Repository for Alzheimer’s Disease (NCRAD), which receives government support under a cooperative agreement grant (U24 AG21886) awarded by the National Institute on Aging (NIA), were used in this study. We thank contributors who collected samples used in this study, as well as patients and their families, whose help and participation made this work possible.

Data for this study were prepared, archived, and distributed by the National Institute on Aging Alzheimer’s Disease Data Storage Site (NIAGADS) at the University of Pennsylvania (U24-AG041689-01)

